# Heterogenous adhesion of follower cells affects tension profile and velocity of leader cells in primary keratocyte collective cell migration

**DOI:** 10.1101/2020.12.08.417063

**Authors:** Baishali Mukherjee, Madhura Chakraborty, Arikta Biswas, Rajesh Kumble Nayak, Bidisha Sinha

## Abstract

Single cell studies demonstrate membrane tension to be a central regulator of lamellipodia-driven motility bringing in front-coherence. During collective cell migration, however, tension mapping or existence of intracellular tension-gradients and the effect of cell-cell interactions have remained unexplored. In this study of membrane fluctuations and fluctuation-tension of migrating primary keratocyte cell-sheets, we first show that some leader cells are followed by followers which remain de-adhered from the substrate while being attached to other cells and thus appear to be “taking a ride”. A subtle yet significant enhanced long-timescale velocity in these leaders indicate increased directionality. Intriguingly, such leaders mostly have front-high tension gradients like single keratocytes, while followers and other leaders usually display front-low membrane tension gradients. The front-high tension gradient and higher membrane tension observed in these leaders, despite the high cell-to-cell variability in membrane tension demonstrate how leader-follower interactions and heterogenous adhesion profiles are key in collective cell migration.

## Introduction

Collective cell migration is central to processes like wound healing, development, and metastasis of cancerous cells. Mechanisms of motility vary from system to system (Ananthakrishnan and Ehrlicher, 2007), however, in single cells that use lamellipodia-driven motility, adhesion and membrane tension play crucial roles. The front-rear polarity is established by lipids and proteins implicated in responses to chemotactic signals; directed vesicle trafficking; regulation of actin nucleation, polymerization, filament-organization; and the regulation of integrin-based adhesion (Ridley et al., 2003). Such processes together impact the cell mechanically leading to coordinated movement of the front and retraction of the rear. As an outcome, not only is the tension of the plasma membrane and its distribution affected, tension also integrates the different inputs and forms the central mechanical regulator of motility. In cells like nematode sperm cells (Batchelder et al., 2011) and neutrophils (Houk et al., 2012), high tension enhances directional persistence while in isolated single keratocytes (Lieber et al., 2015), tension has a front-to-back gradient within the cell with high tension at the front. Optical-trap based experiments (Keren et al., 2008; Lieber et al., 2015) as well as theoretical modelling (Lin et al., 2018) attribute these intracellular tension-gradients to the continuous actin polymerization forces at the front and myosin-based contractile forces at the rear. How does the mechanical environment matter? In studies on keratocytes, interestingly, the actomyosin distribution was found to be less sharp in those keratocytes which were not strictly isolated but tethered to the lagging cell-sheet (Svitkina et al., 1997). Moreover, even in single A2780 cells undergoing durotaxis (Hetmanski et al., 2019), the reported front-high tension gradient was lost when the substrate is uniformly rigid. Both examples underscore how differences in interactions felt at the front and the back of single cells affect its directionality and membrane tension gradient.

Like in single cell motility, collective cell migration also involves players like: actin, Myosin II, integrins, proteolytic agents, cell-cell interactions and cell-matrix interactions among others (Ilina and Friedl, 2009). However, in contrast to the extensive studies about membrane mechanics in single cells, tension measurements have not been done in cell sheets displaying collective cell migration, to the best of our knowledge. How keratocytes that lead the cell sheets (leaders) and cells following leaders (followers) differ in their tension profiles from single isolated cells and how leader-follower interactions impact efficiency of collective migration, therefore, need thorough investigation.

We employ Interference Reflection microscopy (IRM (Biswas et al., 2017; Limozin and Sengupta, 2009)) to image single cells in keratocyte cell sheets and quantify the temporal fluctuations in membrane-height of the basal plasma membrane from the glass-coverslips. These primary keratocytes emerge out of fish scales and move together as a front. Fluctuations are further used to derive effective “fluctuation tension” (Shiba et al., 2016) of the membrane – a quantity that matches the “mechanical frame tension” for over 5 decades of tension values (Fournier and Barbetta, 2008). It is to be noted that henceforth in this paper, the usage of the term “membrane tension” pertain to measurements reported herein, would imply the fluctuation tension of the plasma membrane. Using this non-invasive method, we report details of adhesion to substrate and are able to not only measure but also map membrane tension during collective cell migration.

## Results and Discussions

### Banded adhesion pattern in collective cell sheets of keratocytes

Cell sheets emerging out of fish scales within an hour of incubation, keep expanding in a fan-like shape (**Movie 1**, **Fig. 1A**) with an edge velocity of ~ 0.98 μm/sec (**Fig. 1B**). The whole front line of the sheet is composed of clear “leaders” (**Fig. S1A**). While the sheet appears to be continuous in phase-contrast and DIC imaging, the adhesion status of cells can be best assayed by IRM. IRM shows adhered regions as dark pixels; as distance of the membrane keeps increasing from the glass coverslip, the intensity increases till a maxima is reached at ~100 nm, beyond which intensities start dropping again. Calibration with objects of known topology (~ 60 μm diameter beads) is used for quantifying the conversion of intensity to relative height (distance of basal membrane from coverslip) (Biswas et al., 2017). Leaders show a dark footprint in IRM – indicating their strong although heterogenous attachment to the substrate (**Fig. 1C**). The followers display a previously unknown pattern of adhesion. Most followers are well adhered to the substrate but for some, the cell is visible in DIC, but the attachment pattern is missing in IRM (**Fig. 1C**). This indicates that the cell is more than a micron away from the coverslip (no strong IRM signal) – but attached to its neighbors (visible in DIC). Such cells therefore appear to be “taking a ride”. Such a heterogenous adhesion pattern is not observed in a monolayer of HeLa cells when wounded and left to repair (**Fig. 1D**).

**Figure 1.**
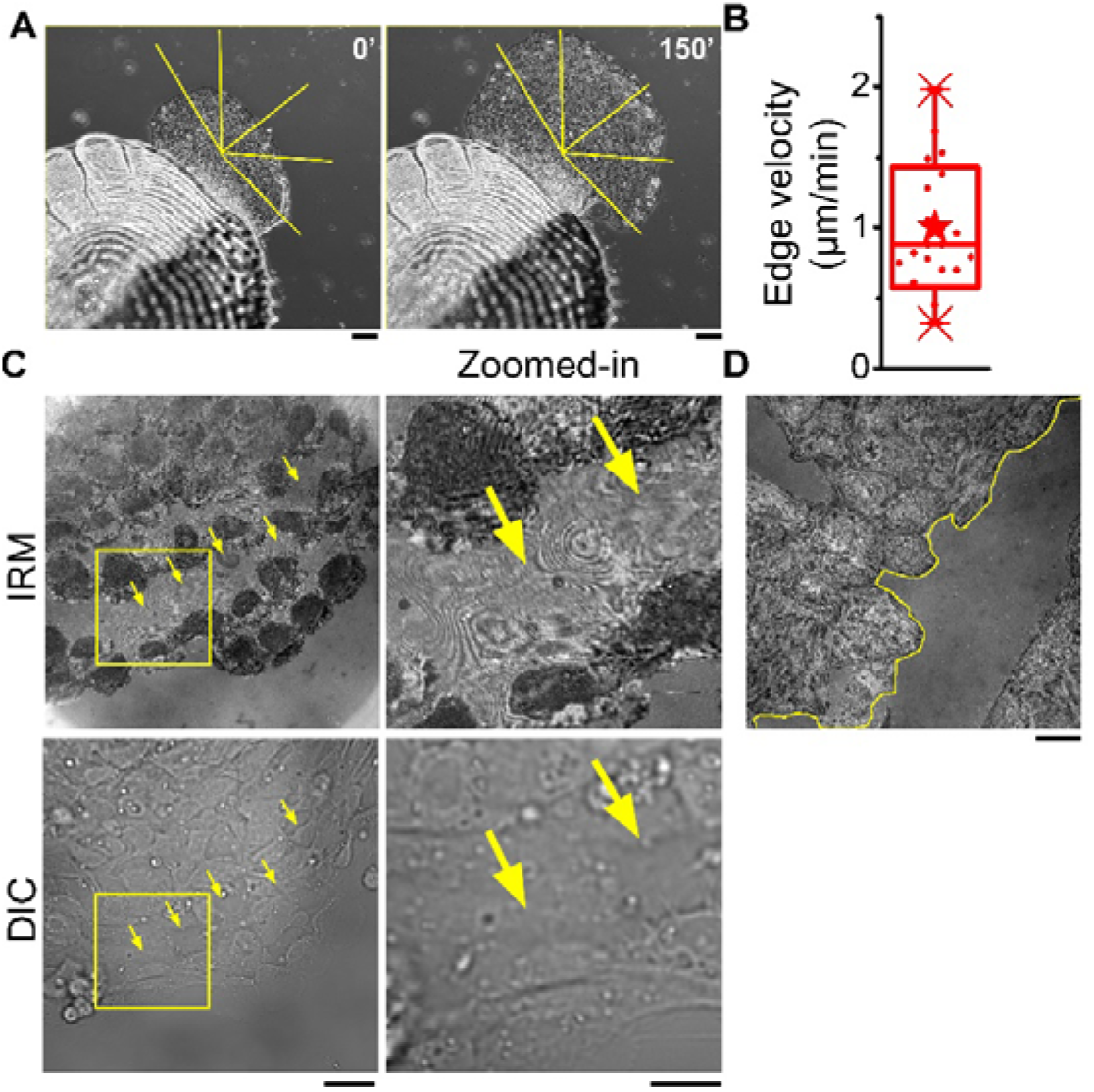
Adhesion profile of keratocyte cell sheets from fish scales and HeLa monolayer. A) Representative images of increase in size of keratocyte cell sheet emerging out of fish scales. Scale bar, 100 μm. B) Velocity of expansion of the cell sheet as shown in A. C) Left: IRM and corresponding DIC images of an edge of a cell sheet. Right: Zoomed-in views of an internal region as marked out in dashed yellow rectangle in C, left. Arrows indicate cells that show fringes in IRM but similar pattern as others in DIC. D) IRM image of a HeLa monolayer with a wound marked in yellow dashed line. Scale bars, 10 μm.

This observation demonstrates that primary cell sheets need not be strictly flat & well adhered, and hence cannot be properly mimicked by HeLa monolayers. While the monolayers are created by first ensuring attachment and then allowing growth, collective cell migration may involve movement into new spaces with varied stiffness and adhesivity. We next investigate if having adhered (termed “sticky”) or de-adhered (termed “non-sticky”) followers affect the movement of leader cells.

### Velocity difference between leaders with sticky vs. non-sticky followers

Velocity of cells were obtained from IRM images by tracking the centroid of the cells obtained from their outlines (**Fig. 2A, B**) either every 41 sec or every 10 min or after 50 min (**Fig. 2C**). Based on the nature of attachment with the followers, leaders were classified as either Leader_S (followed by sticky or substrate-adhered followers) or Leader_N (followed by non-sticky followers). We report that when velocity was measured by comparing centroids over 41sec or 10 min intervals, no significant difference was found between the two types of leaders. However, in the 50 min analysis, the Leader_N pool show a slight (18%) but significantly higher velocity (**Fig. 2C**). Trajectory plots reveal the tortuous trajectory adapted by the Leader_S pool while a relatively straighter path is chosen by Leader_N cells (**Fig. 2D**). This implies that leaders pulled by non-sticky followers perhaps are provided extra cues to retain their directionality. We also observed fluctuations in the spread areas of these individual cells but found no significant difference between the two leader-types (**Fig. 2E**). We next foray to measure membrane tension gradients in these two kinds of leaders.

**Figure 2.**
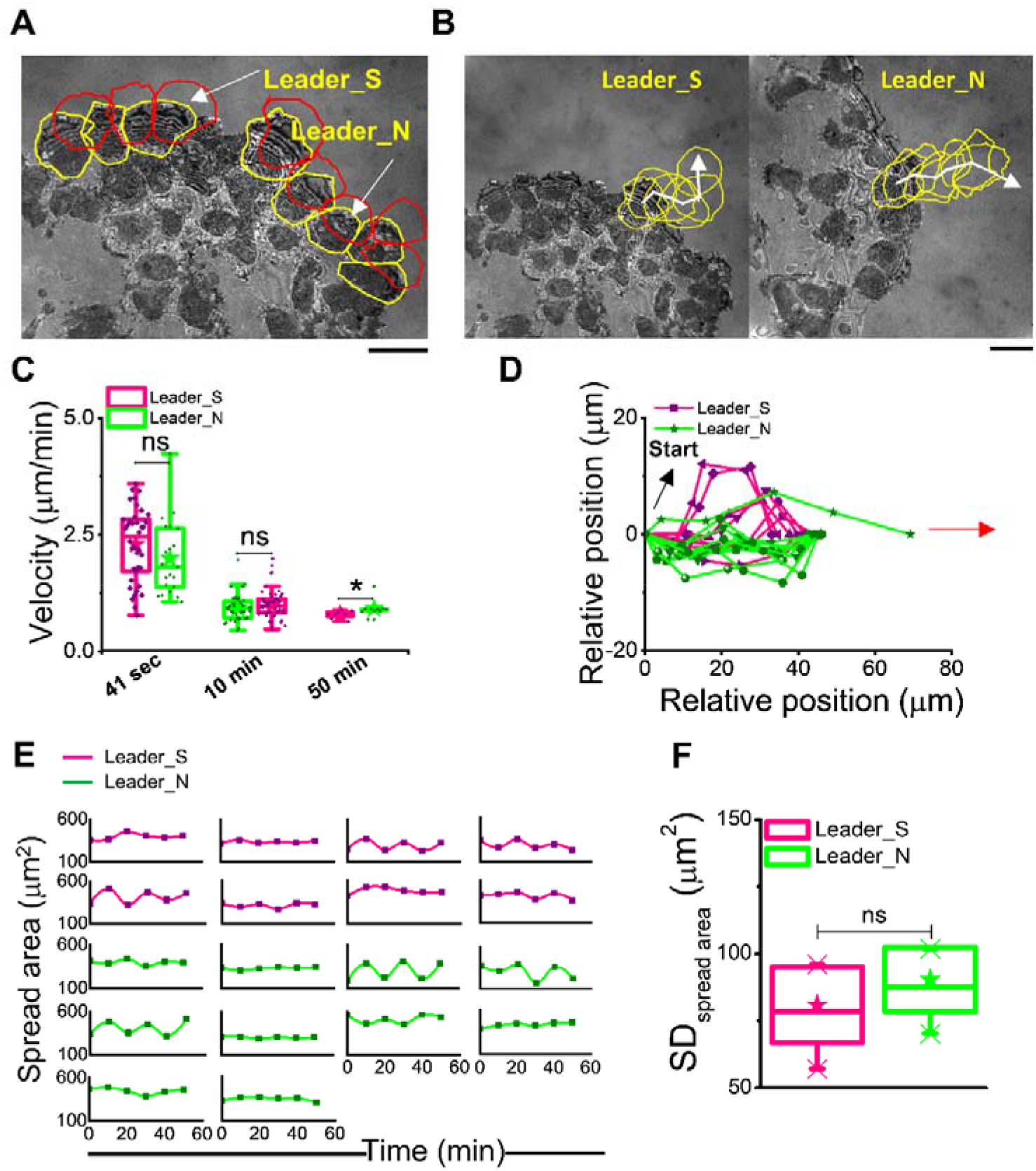
Classifying leaders. A) Representative image of an edge of a cell sheet with yellow lines overlaid around leader cells whose immediate followers are well adhered to the substrate. Red lines outline positions of same cells after 10 min. B) Outlining typical mobility of the two kinds of leader cells every 10 min for 50 min. C) Velocity calculated from time intervals of 41 sec (left), 10 min (centre) and 50 min (right). N_cells_= 59 (“41sec”) (Leader_S= 38 Leader_N= 21), 18 (“10 min”) and 18 (“50 min”) (Leader_S= 8, Leader_N= 10). D) Trajectory plots of Leader_S and Leader_N cells (x vs. y coordinate at different time points) when translated to (0,0) and rotated such that vector connecting first and last points falls on the x-axis, together with black and red arrows denoting start point (0,0) and direction of cell migration respectively. E) Fluctuations in cell spread area of individual cells during migration. F) Comparison of standard deviation of spread areas at different time points together between two leaders’ population. Statistical analysis was performed using Mann-Whitney U test. ns denotes p value > 0.05. * denotes p value < 0.05. Scale bars: 10 μm.

### Fluctuations and Tension ratio of front to mid part of single leader cells with sticky or non-sticky followers

Time-lapse IRM imaging (at 50 frames per sec for 2048 frames) reveals the dynamics of the adhesion profile, with bigger patterns remaining unchanged while smaller fluctuations clearly captured (**Movie 2**). After calibration (see Methods), amplitude of the temporal height fluctuations, SD_time_, is calculated from the time-lapse images and averaged over 6×6 pixel regions (~432 nm × 432 nm) which are specially selected (see Methods) and termed First Branch Regions (FBR). These regions are termed so because they are identified to fall in the first ~100 nm of the coverslip corresponding to the first branch of expected Intensity-height pattern of the interference created, in which the calibration used is applicable (Biswas et al., 2017). Effective tension is calculated from the power spectral density (PSD) of the fluctuations by considering contributions of the membrane’s tension, confinement by the cytoskeleton/substrate, bending rigidity, effective temperature and the effective fluid viscosity around the membrane (Biswas et al., 2017; Biswas et al., 2019).

Fluctuations are evaluated at the front of single cells covering 5 μm from the front edge and the middle section covering the next 5 μm (**Fig. 3A**). Tension maps reported from computational modeling of tension gradients (Fogelson and Mogilner, 2014) are used to decide the front-to-mid segregation with width of 5 μm to strike a balance between increasing sampling and constricting to narrow range of tension. Note that the rear of the cell is not reported here due to the presence of the nucleus in this region. We first show IRM images and corresponding tension maps of typical leader cells representing both kinds of tension gradient for each kind (**Fig. 3A, Fig. S2**). Tension maps are not restricted to pixels that are strictly within the usual ~ 100 nm from the coverslip or the first branch (~ 70% of the basal membrane). Hence, although they nicely capture the tension distribution, we perform analysis at FBRs selected rigorously to compare the front and mid regions of both kind of leader cells. Tension maps validate the choice of ~ 5 μm as the width of the regions. The examples shown represent one front-high and one front-low tension gradients each for both the leader-types. Both leader types show enhanced fluctuations at their fronts (**Fig. 3B**). The ratio of front-to-mid SD_time_ when compared with 0 (by z-test): displayed significant increase from 0 (Leader_S p value =0, Leader_N p value = 5E-240) for both the pools. This indicates that temporal membrane flickering is enhanced at cell fronts. We also compare the level of damping of membrane fluctuations, captured by the parameter “exponent” (**Fig. S2D**) whose absolute value reduces as the fluctuations are damped by restrictive surroundings (Brochard and Lennon, 1975; Gov et al., 2003; Biswas et al., 2017).

**Figure 3.**
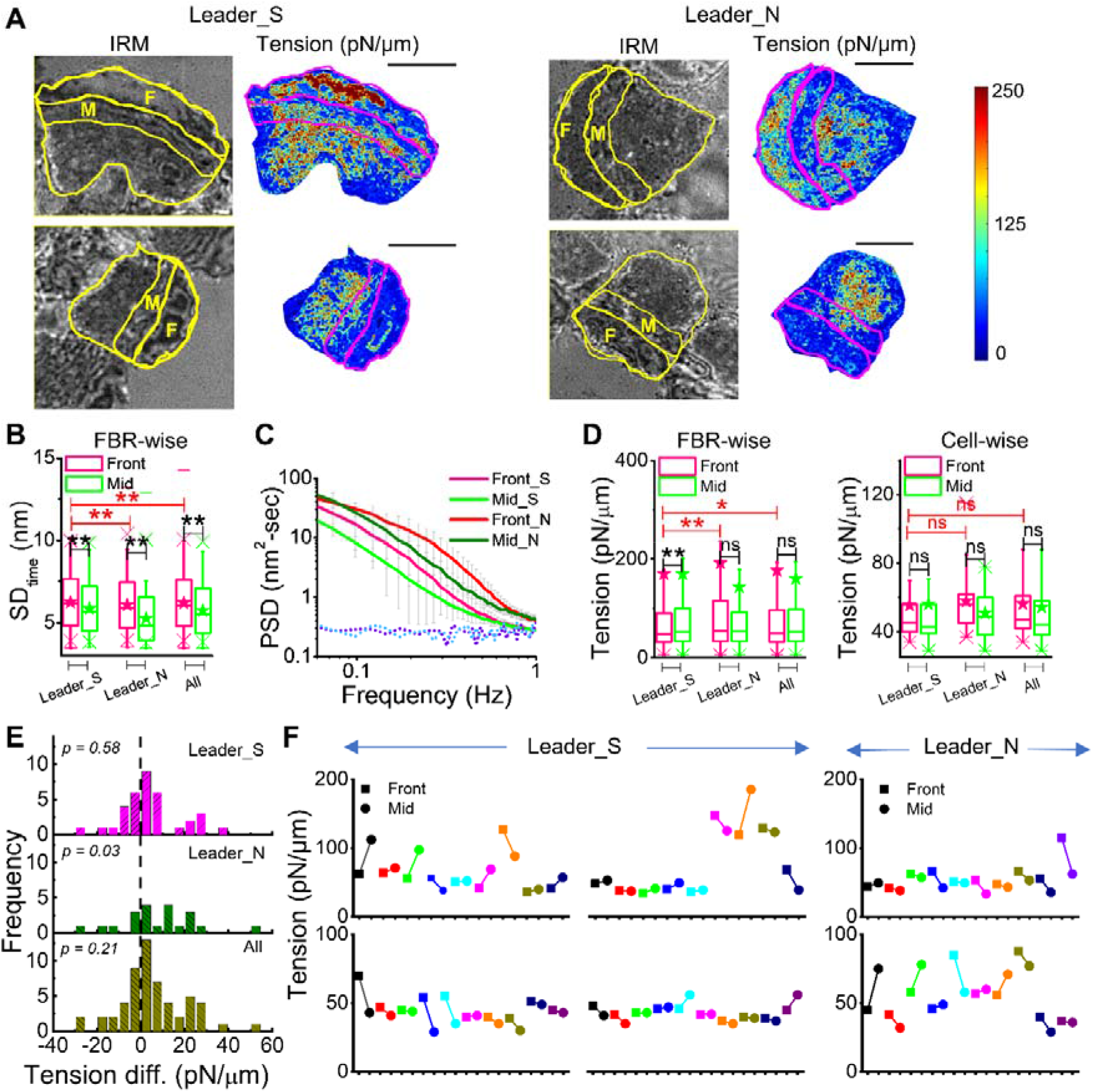
Gradient of fluctuations and tension in leaders. A) Representative IRM images and corresponding tension maps of leader cells (Leader_S and Leader_N) with the front region and the middle region outlined and marked as F and M respectively. B) Comparison of SD_time_ between front and mid FBRs among Leader_S, Leader_N and all leaders. N_Front-FBRs_= 13444 N_Mid-FBRs_= 10271; N_cells_= 38; N_Front-FBRs_= 4025 N_Mid-FBRs_=3039; N_cells_= 21. C) Average power spectral density (PSD) of front and mid regions of leaders followed by sticky cells (Leader_S) and leaders followed by non-sticky cells (Leader_N); N_cells_ = 4; D) Left: FBR wise comparison of tension of front and mid regions among multiple leaders and clubbed together (Leader_S front= 5115, mid=4444; Leader_N front=2363 mid=2481). Right: Cell wise comparison of tension of front and mid regions among multiple leaders and clubbed together (Leader_S =38, Leader_N=21). E) Distribution of tension difference between front and mid regions among different population of leaders. F) Comparison of tension at front and mid-part of each cell, where lines connect front (square symbols) and mid (circle symbols) of each cell. Statistical analysis was performed using Mann-Whitney U test. ns p value > 0.05, * p value< 0.05, ** p value< 0.001. Scale bars: 10 μm.

The PSDs (**Fig. 3C**) reveal differences between front and mid regions which are further fit with models (see Methods) to understand the mechanical parameters that drive the changes (**Fig. 3D, S2**). Pooling all FBRs obtained from fronts of Leader_N cells separately from the FBRs from mid-region showed no difference in tension (**Fig. 3D**). Leader_S, however, showed higher tension in mid-region when all FBRs from all cells were compared with those from mid (**Fig. 3D**). However, when each cell’s front FBRs are averaged and compared with that particular cell’s mid-section, and the difference in tension is pooled from all cells, Leader_N shows significantly higher tension at the front (**Fig. 3E**). In single such cells, the tension difference is found to be non-significant (**Fig. 3E**). Interestingly the front regions of Leader_N has higher tension than front regions of Leader_S (**Fig. 3D**, red star), corroborating the role of tension since high tension has been previously attributed to aid directional motion as observed in Leader_N cells (**Fig. 2**) in this study. This is true even when front as well mid sections are pooled together for comparison (**Fig. S2**). We plot the front-mid tension values for every single cell to reveal the level of heterogeneity (**Fig. 3F**). 16/38 (42%) Leader_S cells show front-low tension profile while 7/21 (33%) Leader_N cells show front-high tension gradient.

Together we demonstrate how the spatial tension profile in leaders can appreciably vary between cells from the reported single cell profiles and can be regulated by the leader-follower interactions bringing in asymmetry in the forces felt. Despite the cell-to-cell variability in tension gradients, the data so far strongly suggests that a front-high tension gradient is advantageous in maintaining directionality of leaders during collective cell migration in keratocytes.

Do followers or cells not facing the edge but in inner layers have similarly heterogenous tension gradients?

### Followers usually display front-low tension gradient

Followers in the second line of cells (from the front) show front-low tension profile irrespective of the tension profile of the cell leading them (**Fig. 4A**). Followers in the third line of cells also show front-low tension profile (**Fig. 4B**) even if the cells following the followers are substrate-detached (**Fig. 4C**). We also show that for the 50 min period over which the sheet was followed; no switching of leader states could be observed. Leaders with non-sticky followers retained their identity as they moved (**Fig. 4D**). The maintenance of directionality hence seems to be the main advantage of front-high tension gradient. For inner cells, the data suggests, the cell’s front is usually maintained at lower tension.

**Figure 4.**
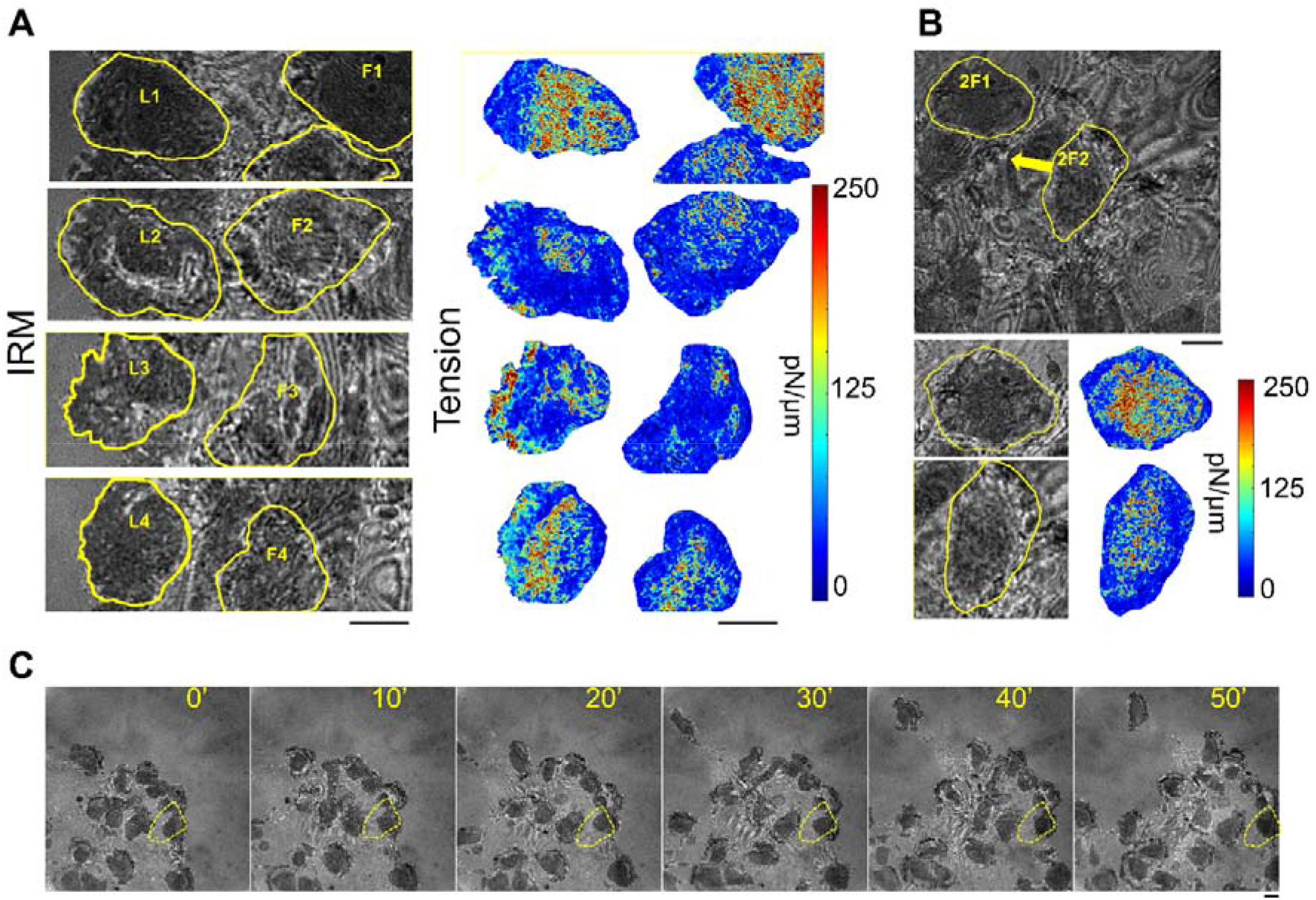
Tension enhanced in deeper layers. A) Left: Representative IRM images of edge of a cell sheet with yellow outlines of leaders marked as L1-L4 and 1^st^ layer of followers marked as F1-F4. Right: tension maps of the same. B) Top: IRM image of two 2^nd^ layers of followers marked as 2F1 and 2F2 and the direction of migration denoted by yellow arrows. Bottom: corresponding tension maps showing enhanced membrane tension in mid and rear portion. C) Representative images of a cell edge followed for 50 min with yellow lines outline a follower that remains “sticky” for the complete period. Scale bars: 10 μm, statistical analysis was performed using Mann-Whitney U test. ns p value > 0.05, ** p value< 0.001. Scale bars: 10 μm.

We therefore conclude through this study on primary cells with non-invasive techniques that that owing to the different inputs from the surroundings, single cells show heterogeneous tension profiles during collective cell migration. Despite this, tension gradient exist and a front-high tension gradient is linked to cues from the rear and demonstrated to still induce directionality in movement. It is the first such study revealing intracellular tension gradients in leaders and followers during collective cell migration. The observation of cells “taking a ride” is unique and could be resolved due to IRM’s is high sensitivity to distance of the basal membrane from the coverslip. Through this study we have been able to demonstrate the strength of IRM in tension mapping. We also show that the keratocyte cell-sheet system offers a new dimension of collective cell migration where cell sheets may adapt other geometries than the strictly 2D geometry usually studied.

## Material and Methods

### Fish and scale collection

Primary keratocyte cultures were collected from the Malawi golden cichlid (*Melanochromis auratus)* as per established protocols (Svitkina et al., 1997; Rapanan et al., 2014 Sun et al., 2016). Briefly, a fish was first anaesthetized with a 10% ethyl p-aminobenzoate for 3-5 min, before 4-5 scales were extracted. Scales were placed on glass-bottomed Petri dish with the inner side of the scale facing the surface. Small metal nuts placed on top of each scale were used to aid the attachment. Within 5 min, 2 ml of growth media was added in the dish and incubated at lab temperature (around 23 °C) for 3 hr. Growth media consisted of Leibovitz-15 (L-15) media (Gibco, Thermo Fisher Scientific, US) supplemented with 15 mM HEPES buffer (Sigma- Aldrich), 10% FBS (Fetal Bovine Serum) (Gibco, Thermo Fisher Scientific) and 1% Anti-anti (Antibacterial-antimicrobial) (Thermo Fisher Scientific). The final pH adjusted to around 7.3 (Sun et al. 2016).

### Imaging

Olympus IX81 microscope (Olympus, Japan) equipped with CMOS camera was used for capturing DIC/Phase contrast images of cell sheet at different magnifications (10x, 40x, 60x) for visualizing migration.

For IRM imaging, Nikon Eclipse Ti-E motorized inverted microscope (Nikon, Japan) was used. Cells were imaged under 60x Plan-Apo (water immersion, NA 1.22) with an external 1.5x magnification objective with a CMOS camera (ORCA-Flash 4.0, Hamamatsu, Japan). A 100W mercury arc lamp, an (546 ± 12 nm) interference filter and a 50-50 beam splitter were used, and fast time-lapse imaging was performed at 50 frames/sec for 2048 frames.

### Calibration for IRM

For converting IRM image to membrane topology (height of basal membrane from glass coverslip) four steps were undertaken. First, stuck 60 μm diameter polystyrene beads (Bangs Laboratories) were imaged at various exposure times. Secondly, the radial intensity profiles were used to calculate the conversion factor for intensity difference (ΔI, au) to height (Δh, nm) conversion for the whole range of contrast obtained in the images. Contrast of an interference image is characterized by the maximum and the minimum or background intensity. Altering contrast mimics the effect of altering reflectivity. For a particular conversion factor to be applicable on cells, the regions must fall in the first branch with a membrane-height ranging from 0-100 nm and the contrast of the interference image must match with the bead-image used to calculate the factor. The third step thus involved measuring the contrast of the images of cells and using the interpolated bead-derived data to estimate the conversion factor for the particular contrast. Finally, the conversion factor was applied to first branch pixels which were selected by checking if they could be spatially connected to a first minima without passing through a first maxima (Biswas et al. 2017).

### Analysis of edge and cell velocity

Sheet velocity was calculated by first getting kymographs of cell sheets across radial lines and extracting edge velocity as the distance covered by the expanding edge per unit time. Cell velocities were calculated by drawing the outline of the cells, finding centroids and finding distance between centroids per unit time.

### Analysis of fluctuations

The first part of the analysis characterizes the amplitude of the temporal fluctuations at particular pixels and termed SD_time_. It is the average standard deviation of the relative height at any particular pixel with the averaging done over a 432 × 432 nm^2^ region comprising of 6×6 pixels. The second part of the analysis characterizes the distribution of the fluctuation amplitude (or power) across different frequencies. The power-spectral density (PSD) of fluctuation signal is plotted as log (PSD) versus log (Frequency). The slope between 0.04 – 0.4 Hz of the log(PSD)-log(frequency) plot is fitted to a straight line, and defined as exponent. All the mechanical parameters were derived from fitting the PSDs of FBRs to an equation:

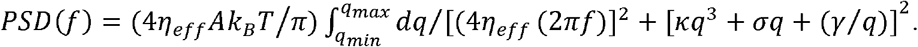

The value of bending rigidity (κ) here fixed as 15 k_B_T. Here □_eff_ denotes effective cytoplasmic viscosity, A denotes active temperature, σ is membrane tension and γ is confinement.

## Author contributions

BS conceptualized the project. BM setup the model system. BM and MC performed the experiments. MC analyzed the experiments. AB and RKN mapped the parameters. BS, MC, and AB wrote the paper and made the display items. BS acquired the funding for the work. All authors edited the paper.

## Acknowledgements

BS acknowledges support from Wellcome Trust/DBT India Alliance fellowship (grant number IA/I/13/1/500885) and SERB [grant number SERB_CRG_2458]. The authors are grateful to DST-Inspire, CSIR and IISER Kolkata for providing scholarship to BM, MC, and AB, respectively. The authors are also thankful to Dr. Anuradha Bhat and her team (IISER Kolkata) for helping with the initial maintenance and care of the fishes, Mr. Ritabrata Ghosh & IISER Central Imaging Facility (CIF) for brightfield imaging, and DIRAC supercomputing facility for tension mapping.

