## Supplementary Material for "Heterogenous adhesion of follower cells affects tension profile and velocity of leader cells in primary keratocyte collective cell migration"

**Figure S1**

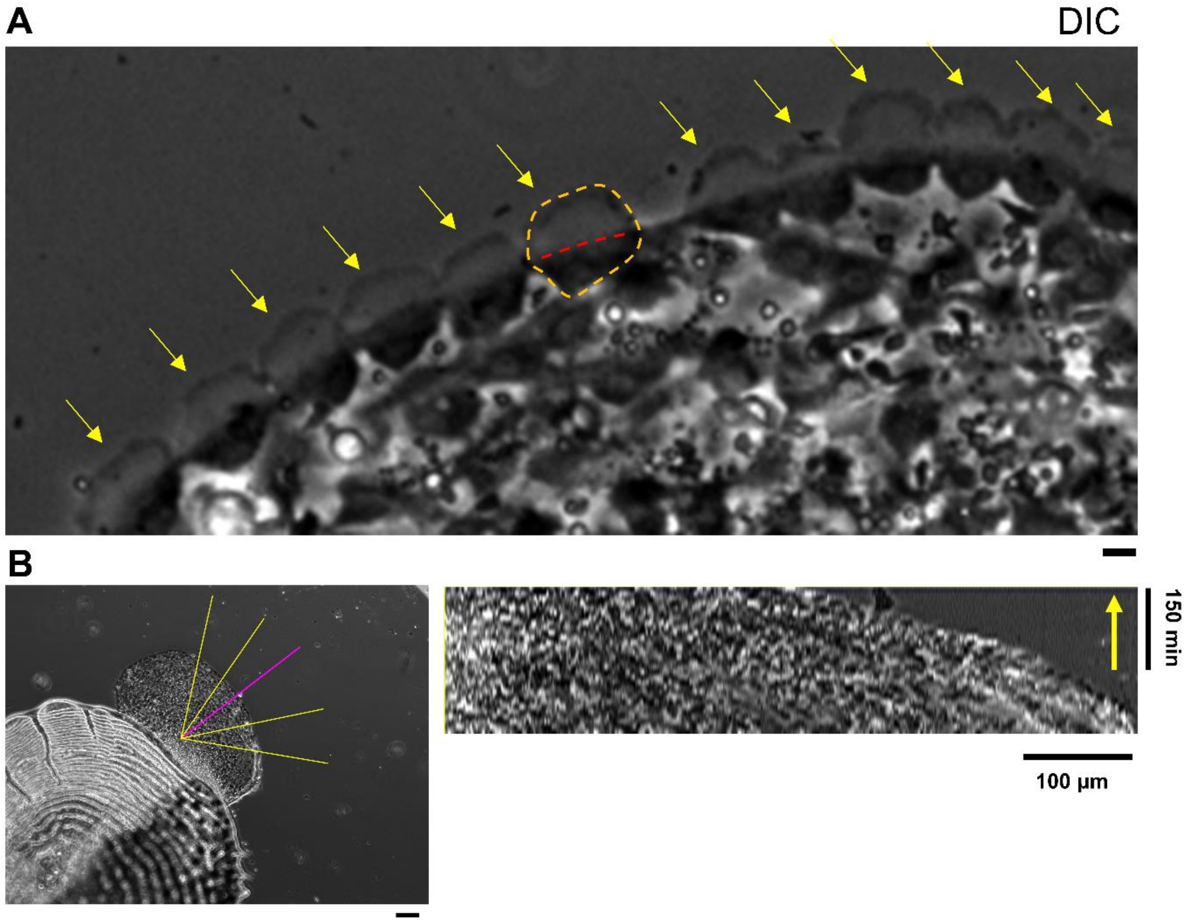


***Figure S1: Edge velocity calculation*** *A) Representative DIC image of a typical keratocyte sheet showing all the leaders denoted by yellow arrows and a typical line like pattern at the mid region marked by magenta dotted structure. B) Left: Cells moving out from under a scale, with lines in yellow drawn to analyse from kymographs to measure velocity along those ROIs. Right: A typical kymograph (calculated from pink line ROI displayed in B) showing edge velocity of a cell sheet. Scale bar 100 µm. Direction of time showed in yellow arrow.*

### Figure S2

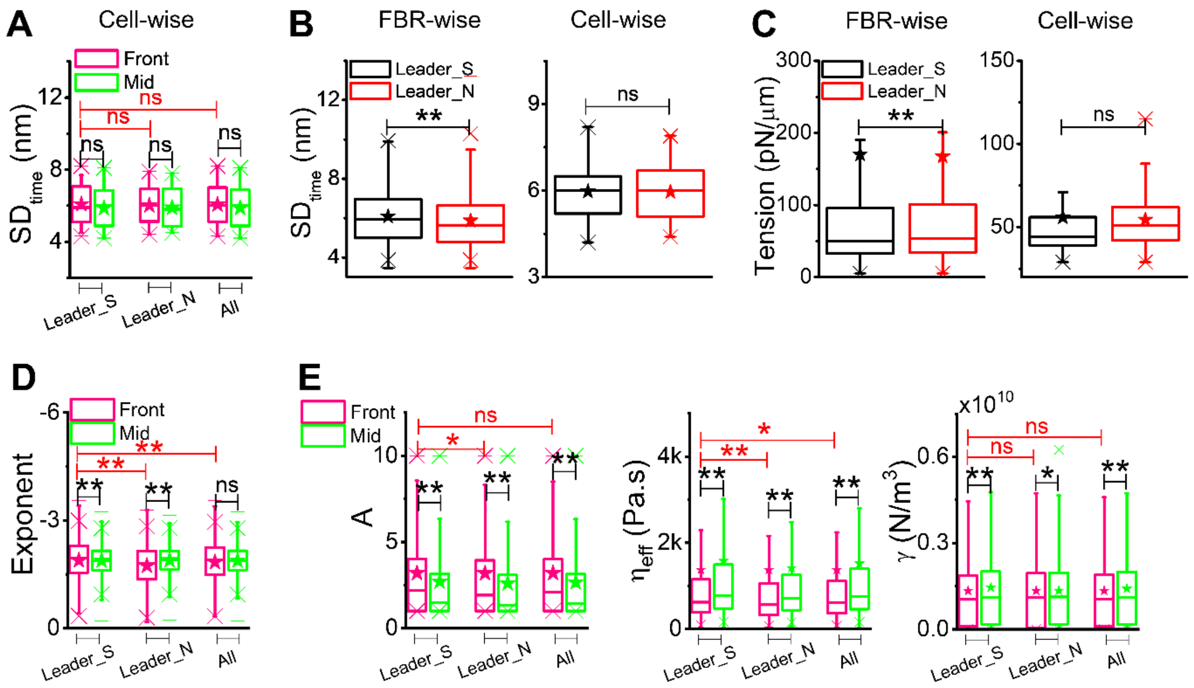


***Figure S2: Box plot comparison of different mechanical parameters*** *A) Cell wise comparison of $SD_{time}$ between front and mid parts of leader cells (Leader_S=38, Leader_N=21, Total=59) B) Left: FBR wise comparison of $SD_{time}$ between Leader_S and Leader_N cells with front and mid clubbed together (Leader_S= 23192, Leader_N=8886), Right: cell wise comparison of $SD_{time}$ with front and mid clubbed together (Leader_S=38, Leader_N=21). C) Left: FBR wise comparison of membrane tension between two different pool of ledaers with front and mid clubbed together (Leader_S=9559, Leader_N=4844). Right: cell wise comparison of the same (Leader_S=38, Leader_N=21). D) FBR wise comparison of exponent between front and mid among different pool of leaders and all leaders clubbed together (Leader_S front=13373, mid=9819, Leader_N front=6580, mid=5345). E) Left: FBR wise comparison of active temperature between front and mid among different pool of leaders and all leaders clubbed together. Centre: FBR wise comparison of effective cytoplasmic viscosity between front and mid among different pool of leaders and all leaders clubbed together. Right: FBR wise comparison of confinement between front and mid among different pool of leaders and all leaders clubbed together (Leader_S front=5115, mid=4444, Leader_N front=2363, mid=2481) Statistical analysis was performed using Mann-Whitney U test. ns denotes p value > 0.05. * denotes p value < 0.05 ** denotes p value <0.001.*

**Figure S3**

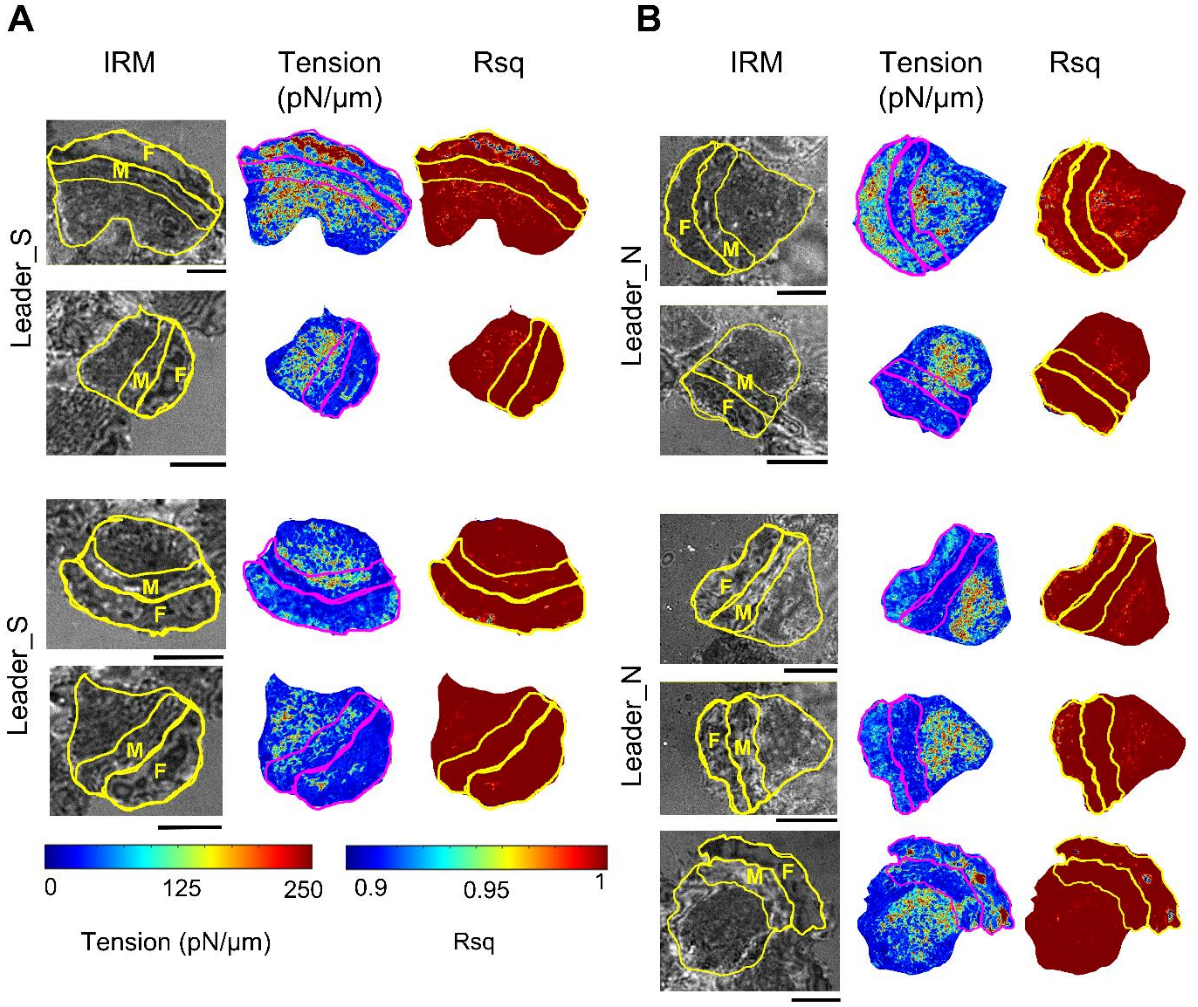


***Figure S3: Tension and their corresponding R-square maps*** *A) Left: Leader_S with front and mid regions marked as F and M. Middle: their respective tension maps, front and mid marked with magenta lines. Right: corresponding R square maps, front and mid marked with yellow lines B) Left: Leader_N with front and mid regions marked as F and M. Middle: respective tension maps, front and mid marked with magenta lines. Right: corresponding R square maps, front and mid marked with yellow lines. Scale bar 10 µm.*

**Figure S4**

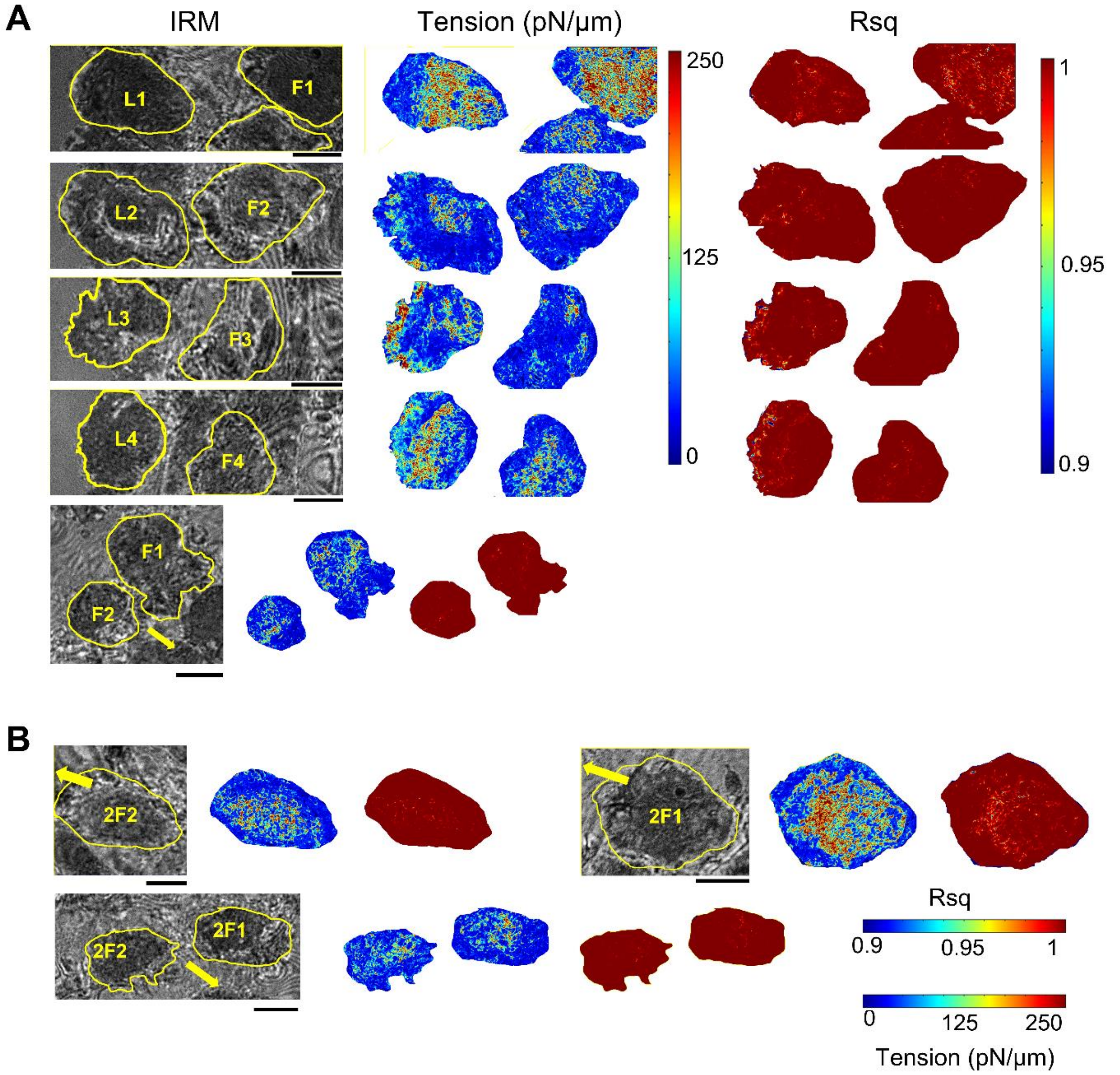


***Figure S4: Tension map and corresponding R-square maps of leaders and followers*** *A) Left: (top to bottom) IRM images of leaders with first layer of followers marked as L1-L4 and F1-F4 respectively. Middle: tension maps of leaders with 1st followers. Right: R-square maps of the same. B) (top and bottom) IRM images of 2nd layer (2F) of followers with their respective tension and R square maps beside them respectively, direction of migration marked with yellow arrows. Scale bar 10 µm.*

**Table S1**

| Figure 1B | | | | | | | | |
|---|---|---|---|---|---|---|---|---|
| **Parameters** | **Condition** | **N** | **n** | **Mean** | **SD** | **SEM** | **Median** | |
| Edge velocity (µm/min) | Growth media | 4 | 20 | 1 | 0.4 | 0.1 | 0.9 | |
| **Figure 2C** | | | | | | | | |
| **Parameters** | **Condition** | **N** | **n** | **Mean** | **SD** | **SEM** | **Median** | **p-values (wrt Leader_S)** |
| Velocity (µm/min) | Leader_S (41 sec) | 38 | 38 | 2.3 | 0.79 | 0.13 | 2.55 | |
| | Leader_N (41 sec) | 21 | 21 | 1.94 | 0.77 | 0.17 | 1.8 | 0.10805 |
| | Leader_S (10 min) | 8 | 40 | 0.93 | 0.29 | 0.05 | 0.91 | |
| | Leader_N (10 min) | 10 | 50 | 1 | 0.3 | 0.04 | 0.97 | 0.24391 |
| | Leader_S (50 min) | 8 | 8 | 0.78 | 0.09 | 0.03 | 0.79 | |
| | Leader_N (50 min) | 10 | 10 | 0.92 | 0.18 | 0.06 | 0.9 | 0.01133 |
| **Figure 2F** | | | | | | | | |
| **Parameters** | **Condition** | **N** | **n** | **Mean** | **SD** | **SEM** | **Median** | **p-values (wrt Leader_S)** |
| $SD_{Spread\ area}$ ($\mu m^2$) | Leader_S | 6 | 8 | 80.91 | 14.12 | 5.76 | 80.85 | |
| | Leader_N | 6 | 10 | 90.47 | 11.92 | 4.87 | 92.32 | 0.17 |
| **Figure 3B** | | | | | | | | |
| **Parameters** | **Condition** | **N** | **n** | **Mean** | **SD** | **SEM** | **Median** | **p-values (wrt Front)** |
| $SD_{time}$ (nm) | Front; Leader_S | 38 | 13373 | 6.2 | 1.4 | 0.01 | 6.1 | |
| | Mid; Leader_S | 38 | 9819 | 5.9 | 1.3 | 0.01 | 5.7 | 2E-23 |
| | Front; Leader_N | 21 | 6580 | 6.1 | 1.4 | 0.01 | 5.9 | |
| | Mid; Leader_N | 21 | 2306 | 5.3 | 1.3 | 0.03 | 4.8 | 3E-16 |
| | Front; All | 59 | 19953 | 6.2 | 1.4 | 0.01 | 6.1 | |
| | Mid; All | 59 | 12125 | 5.7 | 1.4 | 0.01 | 5.5 | 3E-21 |

| Figure 3D | | | | | | | | |
|---|---|---|---|---|---|---|---|---|
| **Parameters** | **Condition** | **N** | **n** | **Mean** | **SD** | **SEM** | **Median** | **p-values (wrt Front)** |
| Tension (pN/µm) | Front; Leader_S | 38 | 5115 | 170 | 527.1 | 7.4 | 47.4 | |
| | Mid; Leader_S | 38 | 4444 | 169.8 | 562.1 | 8.4 | 52.2 | 6E-4 |
| | Front; Leader_N | 21 | 2363 | 191.8 | 490.8 | 10.1 | 53.9 | |
| | Mid; Leader_N | 21 | 2481 | 143.6 | 403.6 | 8.1 | 53.4 | 0.0515 |
| | Front; All | 59 | 7478 | 176.9 | 515.9 | 6 | 49.1 | |
| | Mid; All | 59 | 6925 | 160.4 | 511.1 | 6.1 | 52.7 | 0.06068 |
| **Parameters** | **Condition** | **N** | **n** | **Mean** | **SD** | **SEM** | **Median** | **p-values (wrt Front)** |
| Tension (pN/µm) | Front; Leader_S | 38 | | 55.5 | 27.7 | 4.5 | 45.5 | |
| | Mid; Leader_S | 38 | | 56.2 | 32.6 | 5.3 | 43 | 0.46357 |
| | Front; Leader_N | 21 | | 58 | 18.7 | 4.1 | 56 | |
| | Mid; Leader_N | 21 | | 51 | 15.3 | 3.3 | 49 | 0.24182 |
| | Front; All | 59 | | 56.4 | 24.8 | 3.2 | 47 | |
| | Mid; All | 59 | | 54.3 | 27.7 | 3.6 | 44 | 0.19714 |

**Table S1: Descriptive statistics for all parameters plotted in the main figures.**

**Table S2**

**Figure S3**

| Parameters | Condition | N | n | Mean | SD | SEM | Median | p-values (wrt Front) |
|---|---|---|---|---|---|---|---|---|
| $SD_{time}$ (nm) | Front; Leader_S | 38 | 13373 | 6.24 | 1.42 | 0.012 | 6.13 | |
| | Mid; Leader_S | 38 | 9819 | 5.85 | 1.34 | 0.013 | 5.67 | 0 |
| | Front; Leader_N | 21 | 658.0 | 6.07 | 1.37 | 0.016 | 5.90 | |
| | Mid; Leader_N | 21 | 5345 | 5.25 | 1.31 | 0.027 | 4.82 | 0 |
| | Front; All | 59 | 19953 | 6.19 | 1.41 | 0.009 | 6.05 | |
| | Mid; All | 59 | 15164 | 5.73 | 1.36 | 0.012 | 5.50 | 0 |

**Figure S3B**

| Parameters | Condition | N | n | Mean | SD | SEM | Median | p-values (wrt Front) |
|---|---|---|---|---|---|---|---|---|
| Exponent | Front; Leader_S | 38 | 13373 | -1.89 | 0.58 | 0.005 | -1.96 | |
| | Mid; Leader_S | 38 | 9819 | -1.88 | 0.4 | 0.004 | -1.88 | 5E-14 |
| | Front; Leader_N | 21 | 6580 | -1.74 | 0.57 | 0.007 | -1.8 | |
| | Mid; Leader_N | 21 | 5345 | -1.89 | 0.39 | 0.005 | -1.9 | 2E-30 |
| | Front; All | 59 | 19953 | -1.84 | 0.58 | 0.004 | -1.91 | |
| | Mid; All | 59 | 15164 | -1.88 | 0.4 | 0.003 | -1.89 | 0.23452 |

**Figure S3C**

| Parameters | Condition | N | n | Mean | SD | SEM | Median | p-values (wrt Front) |
|---|---|---|---|---|---|---|---|---|
| A | Front; Leader_S | 38 | 5115 | 3.2 | 2.8 | 0.04 | 2.2 | |
| | Mid; Leader_S | 38 | 4444 | 2.7 | 2.6 | 0.04 | 1.5 | 2E-22 |
| | Front; Leader_N | 21 | 2363 | 3.2 | 3 | 0.06 | 1.9 | |
| | Mid; Leader_N | 21 | 2481 | 2.6 | 2.6 | 0.05 | 1.3 | 7E-13 |
| | Front; All | 59 | 7478 | 3.2 | 2.9 | 0.03 | 2.1 | |
| | Mid; All | 59 | 6925 | 2.6 | 2.6 | 0.03 | 1.4 | 4E-21 |

| Parameters | Condition | N | n | Mean | SD | SEM | Median | p-values (wrt Front) |
|---|---|---|---|---|---|---|---|---|
| $\eta_{eff}$ (Pa.s) | Front; Leader_S | 38 | 5115 | 1376.5 | 3237.3 | 45.3 | 622.9 | |
| | Mid; Leader_S | 38 | 4444 | 1581.3 | 2861.8 | 42.9 | 780.6 | 7E-40 |
| | Front; Leader_N | 21 | 2363 | 1372.4 | 3631.5 | 74.7 | 564.5 | |
| | Mid; Leader_N | 21 | 2481 | 1418.2 | 2888.3 | 58 | 715.8 | 1E-23 |
| | Front; All | 59 | 7478 | 1375.2 | 3366.6 | 38.9 | 606.6 | |
| | Mid; All | 59 | 6925 | 1522.9 | 2872.2 | 34.5 | 753 | 9E-59 |
| **Parameters** | **Condition** | **N** | **n** | **Mean** | **SD** | **SEM** | **Median** | **p-values (wrt Front)** |
| $\gamma$ ($N/m^3$) | Front; Leader_S | 38 | 5115 | 1.3E9 | 1.6E9 | 2.2E7 | 1E9 | |
| | Mid; Leader_S | 38 | 4444 | 1.5E9 | 1.6E9 | 2.4E7 | 1.1E9 | 8E-5 |
| | Front; Leader_N | 21 | 2363 | 1.3E9 | 1.5E9 | 3.2E7 | 1.1E9 | |
| | Mid; Leader_N | 21 | 2481 | 1.3E9 | 1.4E9 | 2.7E7 | 1.1E9 | 0.03323 |
| | Front; All | 59 | 7478 | 1.3E9 | 1.6E9 | 1.8E7 | 1.1E9 | |
| | Mid; All | 59 | 6925 | 1.4E9 | 1.6E9 | 1.9E7 | 1.1E9 | 6E-6 |
| **Parameters** | **Condition** | **N** | **n** | **Mean** | **SD** | **SEM** | **Median** | **p-values (wrt Front)** |
| Rsq. | Front; Leader_S | 38 | 5115 | 0.95219 | 0.05013 | 7.00929E-4 | 0.9717 | |
| | Mid; Leader_S | 38 | 4444 | 0.95977 | 0.04563 | 6.84423E-4 | 0.97867 | 5E-15 |
| | Front; Leader_N | 21 | 2363 | 0.94333 | 0.05588 | 0.00117 | 0.96354 | |
| | Mid; Leader_N | 21 | 2481 | 0.95766 | 0.04623 | 9.5704E-4 | 0.97698 | 2E-19 |
| | Front; All | 59 | 7478 | 0.94946 | 0.05213 | 6.06255E-4 | 0.96947 | |
| | Mid; All | 59 | 6925 | 0.95904 | 0.04584 | 5.56845E-4 | 0.9782 | 9E-30 |

**Table S2: Descriptive statistics for all parameters plotted in supplementary figures.**

**Movie 1**

A representative movie of collective cell migration of cells emerging out of a fish scale. Phase-contrast images were acquired every 10 mins.

**Movie 2**

A representative movie of membrane fluctuations of a leader cell captured by IRM at 50 frames per sec acquisition rate.
